# Genome-wide RNA binding analysis of *C9orf72* poly(PR) dipeptides

**DOI:** 10.1101/2022.10.10.511318

**Authors:** Rubika Balendra, Igor Ruiz de los Mozos, Idoia Glaria, Carmelo Milioto, Hana M Odeh, Katherine M Wilson, Agnieszka M Ule, Martina Hallegger, Laura Masino, Stephen Martin, Rickie Patani, James Shorter, Jernej Ule, Adrian M Isaacs

**Author notes:** These authors contributed equally (Rubika Balendra, Igor Ruiz de los Mozos). Correspondence to Professor Adrian Isaacs and Professor Jernej Ule.

## Abstract

An intronic GGGGCC repeat expansion in *C9orf72* is a common genetic cause of amyotrophic lateral sclerosis and frontotemporal dementia. The repeats are transcribed in both sense and antisense directions to generate distinct dipeptide repeat proteins, of which poly(GA), poly(GR) and poly(PR) have been implicated in contributing to neurodegeneration. Poly(PR) binding to RNA may contribute to toxicity, but analysis of poly(PR)-RNA binding on a genome-wide scale has not yet been carried out. We therefore performed crosslinking and immunoprecipitation (CLIP) analysis in human cells to identify the RNA binding sites of poly(PR). We found that poly(PR) binds to nearly 600 RNAs, with the sequence GAAGA enriched at the binding sites. *In vitro* experiments showed that polyGAAGA RNA binds poly(PR) with higher affinity than control RNA and induces phase-separation of poly(PR) into condensates. These data indicate that poly(PR) preferentially binds to polyGAAGA-containing RNAs, which may have physiological consequences.

## Introduction

A hexanucleotide repeat expansion in the *C9orf72* gene is the most common genetic cause of amyotrophic lateral sclerosis (ALS) and frontotemporal dementia (FTD) (Renton et al., 2011, DeJesus-Hernandez et al., 2011). Several mechanisms of toxicity have been implicated as contributing to the disease process (Balendra and Isaacs, 2018). Dipeptide repeat protein (DPRs) produced by repeat-associated non-ATG translation (RAN) are likely to represent an important toxic entity (Mori et al., 2013, Zu et al., 2013, Ash et al., 2013). Five different DPRs are produced: poly(GA), poly(GP), poly(GR), poly(PA) and poly(PR). Of these, the arginine containing DPRs, poly(GR) and poly(PR) are the most toxic in model systems (Moens et al., 2017). The common pathological hallmark identified in the vast majority of sporadic and genetic ALS cases and a large proportion of FTD cases is mislocalisation and aggregation of the RNA- and DNA-binding protein TDP-43. This pathology is also found in *C9orf72* amyotrophic lateral sclerosis and frontotemporal dementia (C9FTD/ALS), and is likely to be downstream of DPR pathology (Balendra and Isaacs, 2018).

Several mechanisms have been attributed to DPR pathology, and include nucleocytoplasmic trafficking dysfunction, DNA damage and translational inhibition. A number of studies have explored the effect of the arginine-containing DPRs on membrane-less organelles, such as stress granules and nucleoli. Interactome studies have confirmed that poly(PR) and poly(GR) bind to proteins enriched in prion-like low complexity domains (LCDs), many of which are RNA-binding proteins (RBPs) and constituents of membrane-less organelles (Lin et al., 2016, Lee et al., 2016, Boeynaems et al., 2017, Moens et al., 2019, Hartmann et al., 2018, Odeh and Shorter, 2020). LCDs in RBPs facilitate the process known as liquid-liquid phase separation (LLPS), by which membrane-less organelles are formed, and this process is promoted by the presence of RNA (Molliex et al., 2015, Murakami et al., 2015, Patel et al., 2015, Protter et al., 2018). Mutations in TDP-43, FUS and hnRNPA1 cause ALS/FTD, and these mutations are often localised within the LCDs of these RBPs. These mutations increase the formation of amyloid like fibrils and disturb LLPS dynamics. Poly(PR) and poly(GR) disrupt the dynamics of LLPS in membrane-less organelles in cells and impair translation (Lee et al., 2016, Boeynaems et al., 2017, White et al., 2019, Moens et al., 2019, Hartmann et al., 2018, Zhang et al., 2018). These arginine rich DPRs can also undergo LLPS themselves in vitro which is dependent on anion charge, and the presence of RNA dose dependently increases LLPS of poly(PR) (Boeynaems et al., 2017, Boeynaems et al., 2019). It is possible that these interactions with RBPs and other LCD-containing proteins are partly mediated by interactions of poly(PR) with RNA. An interactome analysis of poly(GR)80 expressed in human embryonic kidney cells revealed that it interacts with RBPs and ribosomal proteins, including mitochondrial ribosomal proteins (Lopez-Gonzalez et al., 2016). Several interactions were abolished when samples were treated with RNase A, suggesting some were RNA-mediated. Another study demonstrated that poly(PR) interacts with multiple DEAD-box RNA helicases, and that this is dependent on RNA, suggesting RNA mediates the interaction (Suzuki et al., 2018). Poly(PR)20 peptide, when applied exogenously to human astrocyte cells in culture, leads to alterations in splicing of several mRNAs and a change in abundance of mRNAs encoding ribosomal proteins in particular (Kwon et al., 2014), and some of these RNAs are bound directly by poly(PR) (Kanekura et al., 2016).

However, a genome-wide analysis of poly(PR) binding to RNAs in the cellular context has not been investigated yet. To achieve this goal we used improved iCLIP (iiCLIP), which enables quantitative identification of protein–RNA crosslink sites *in vivo* (Lee et al., 2021), to investigate whether the arginine-containing DPR poly(PR) binds to RNA with some sequence specificity in human cells. We show that poly(PR) directly crosslinks to RNA and shows enriched crosslinking on specific transcripts, including ALS-relevant mRNAs such as Neurofilament Medium Chain (*NEFM*) and Nucleolin (*NCL*). We further show that poly(PR) interacts with nanomolar affinity with GAAGA-containing RNA, which also promotes phase separation of poly(PR).

## Results

### Poly(PR) iiCLIP reveals binding to specific RNAs

To investigate genome-wide DPR binding to RNA we established a denaturing purification of DPR-RNA complexes for CLIP based on the previously established approach (Fig 1A-C) (Huppertz et al., 2014, Lee et al., 2021). We expressed doxycycline inducible triple FLAG tagged PR100, GA100, or triple FLAG tag alone in human embryonic kidney cells (HEK293Ts) (Fig 1B). After 24 hours of exogenous expression, we used UV light to crosslink protein-RNA interactions. We subsequently immunoprecipitated the DPR-RNA complexes using the FLAG tag (Fig S1A, S1B). We then employed the iiCLIP protocol to ligate an infrared adaptor for visualisation of the protein-RNA complexes, and extracted and reverse transcribed the RNA to generate cDNA libraries for high-throughput sequencing (Lee et al., 2021). Infrared visualisation of the DPR-RNA complexes showed much stronger signal in the crosslinked PR100-FLAG cells (Fig 2A lanes 3 and 4) compared to the non-crosslinked PR100-FLAG cells (Fig 2A lane 5), the crosslinked GA100-FLAG cells (Fig 2A lanes 6 and 7), and the crosslinked FLAG only cells (Fig 2A lane 9) – and this difference is especially apparent for the diffuse signal that usually represents proteins crosslinked to longer RNA fragments (Fig 2A). This finding indicates that PR100 directly crosslinks to RNA in human cells.

**Figure 1.**
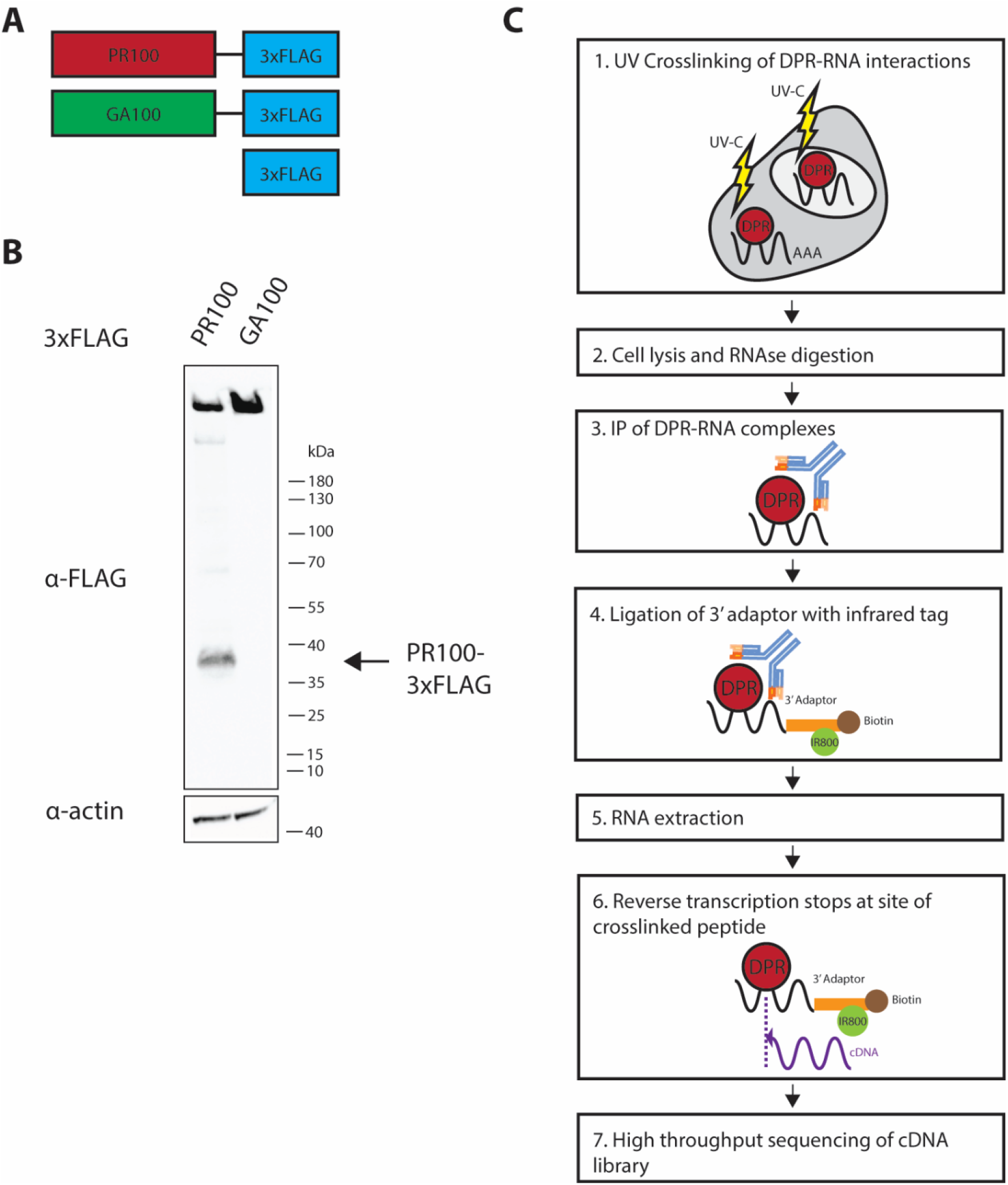
*C9orf72* dipeptide repeat protein iiCLIP pipeline. (A) Diagram of the PR100-3xFLAG, GA100-3xFLAG and 3xFLAG alone constructs used in this study. (B) Anti-FLAG immunoblot 24 hours post-induction of PR100-3xFLAG, GA100-3xFLAG in HEK293T cells. (C) Summary of iiCLIP pipeline for investigation of DPR-RNA direct interaction. Transiently transfected HEK293Ts were UV-crosslinked to stabilise DPR-protein interactions, cells were lysed and digested with RNAse. The FLAG tag was used for immunoprecipitation of DPR-RNA complexes. A preadenylated, infrared dye labelled adaptor was ligated onto the 3’ end of the RNA. RNA was extracted and reverse transcribed, generating cDNA libraries which were high throughput sequenced, and the data analysed to determine sites of binding with nucleotide specificity across the genome.

**Figure 2.**
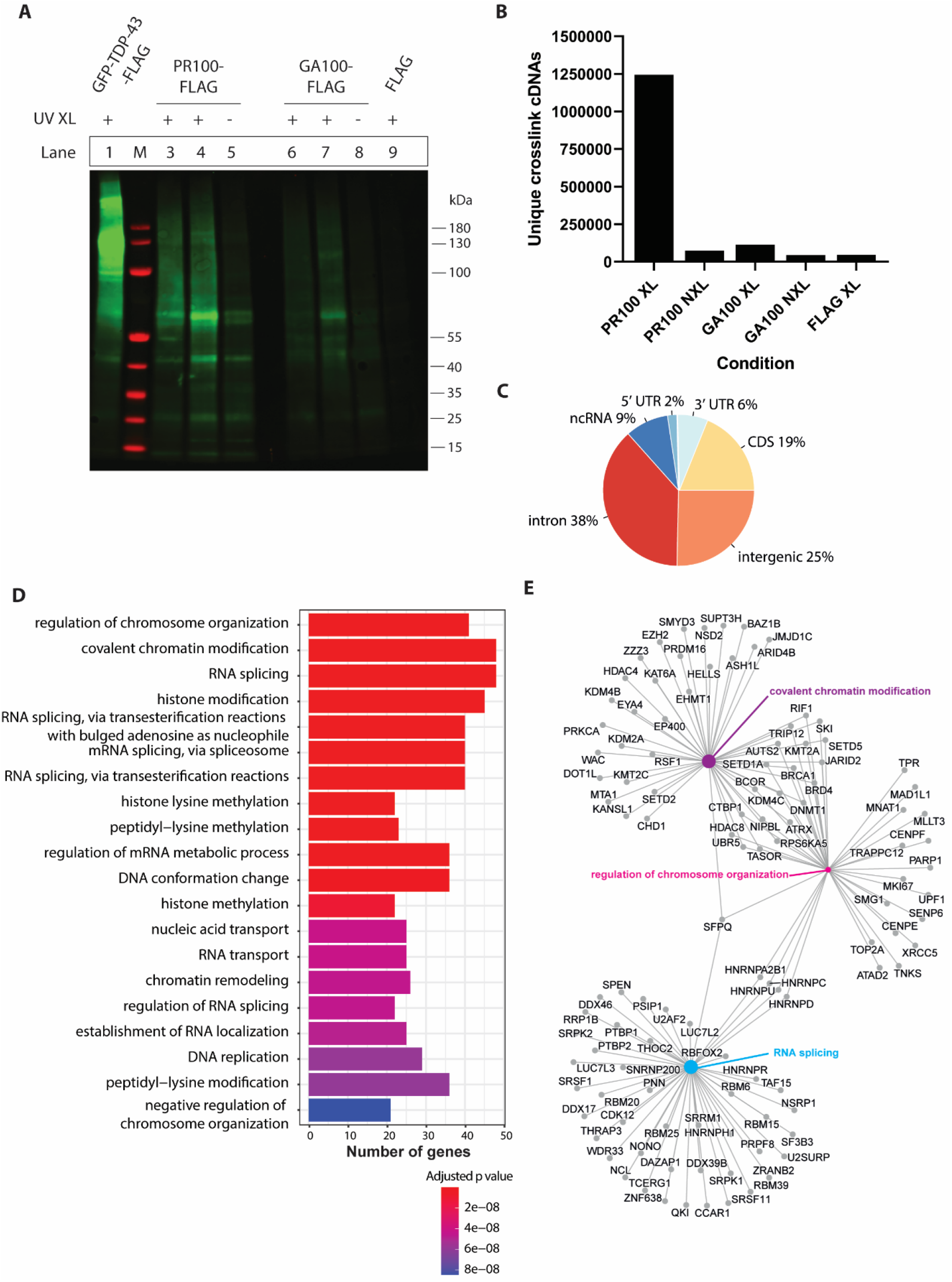
iiCLIP reveals poly(PR) binds to RNA in human cells. (A) Infrared labelled protein-RNA complexes were separated by SDS-PAGE and transferred onto a nitrocellulose membrane. In lane 1, GFP-TDP-43-FLAG was run as a positive control, with a diffuse smear detected above its molecular weight, representing the GFP-TDP-43-FLAG-RNA complexes. Lane M is the protein ladder marker. Lanes 3 and 4 are crosslinked PR100-FLAG cells which have a higher intensity than the PR100-FLAG non-crosslinked cells (lane 5), the GA100-FLAG crosslinked cells (lanes 6 and 7), the GA100-FLAG non-crosslinked cells (lane 8) and the FLAG crosslinked cells (lane 9). The FLAG tag consists of 3XFLAG. (B) The frequency of unique cDNAs which represent individual crosslinking events identified by iiCLIP analysis. (C) Genomic location of PR100 binding sites, in introns, intergenic, coding sequence (CDS), non-coding RNA (ncRNA), 5’ UTR and 3’ UTR, segments. (D) Gene Ontology gene set enrichment analysis of PR100 crosslinked RNAs. Genes of RNAs bound in PR100 samples are represented by their Biological Process. The number of genes from the PR100-FLAG crosslinking dataset in each Gene Ontology category are shown and colour coded by adjusted p value. (E) The genes of RNAs bound in PR100 samples from the top three significant categories within Biological Process (RNA splicing, regulation of chromosome organisation and covalent chromatin modification) are represented in a gene-concept network. The size of the circle for each Biological Process is proportional to the number of genes identified within that category.

Sequencing of the iiCLIP reads revealed over 1,200,000 unique cDNA crosslinking events in the crosslinked PR100-FLAG cells across multiple replicates, with significantly fewer in the control (<74,000) and GA100 (<120,000) conditions (Fig 2B). PR100 crosslink events occurred most frequently in introns, intergenic regions and in coding sequence, with additional signal in non-coding RNAs and 3’ and 5’ UTRs (Fig 2C). We identified 558 mRNAs with high levels of binding (≥200 crosslink events) to PR100 as compared to the controls of PR100 non-crosslinked and FLAG alone samples (Table S1). Examples of genes with the highest numbers of binding events (>1000 crosslink events) included X inactive specific transcript (*XIST*), Metastasis-Related Lung Adenocarcinoma Transcript 1 (*MALAT1), NEFM, NCL*, Nuclear Enriched Abundant Transcript 1 (*NEAT1*) and Heterogeneous Nuclear Ribonucleoprotein U (*HNRNPU*) (Table S1). *XIST* and *NEAT1* were in the top seven genes with the largest number of crosslink events (Table S1), in agreement with their previously identified interaction with poly(PR) through RNA-IP experiments (Suzuki et al., 2019). In addition, the parkin gene (*PRKN*), mutations in which cause Parkinson’s disease (Kitada et al., 1998), had 266 crosslink events. Gene Ontology (GO) enrichment analysis of the 558 mRNAs with the highest number of cross-links to PR100 (Table S1) revealed a significant enrichment involving the biological processes of ‘RNA splicing’, ‘regulation of chromosome organisation’ and ‘covalent chromatin modification’ (Fig 2D and E). There was also significant enrichment in the cellular components of ‘nuclear speckles’, ‘chromosome region’ and ‘centromeric chromosome regions’ (Fig S2A and B), and the molecular functions of ‘ATPase activity’, ‘DNA-dependent ATPase activity’ and ‘helicase activity’ (Fig S2C).

### Poly(PR) binds with high affinity to polyGAAGA RNAs

We analysed enrichment of 5mer motifs in our PR100 iiCLIP dataset, which identified GAAGA as a highly enriched pentameric sequence (Fig 3A), exemplified in the *NCL* and *NEFM* transcripts (Fig 3C and D). UAUAA was a less represented motif (in the bottom 25% of all motifs) surrounding the crosslinking site (Fig 3B). To determine whether there was a differential affinity of poly(PR) for these 5mer RNA sequences, we used biolayer interferometry to compare the affinity of purified PR20 and GP20 peptides to biotinylated RNA oligonucleotides containing five repeats of GAAGA or UAUAA (Table 1). PR20 had stronger affinity for the polyGAAGA RNA with an apparent Kd of 2 ± 0.7 nM (Fig 4A and C) compared to the polyUAUAA RNA, with an apparent Kd of 27 ± 6 nM (Fig 4B and D). There was no evidence of interaction between the DPR GP20 and the polyGAAGA RNA, even at 100-fold higher concentrations (3 – 25 μM for GP20 compared to up to 250 nM for PR20) (Fig 4E). These experiments show that poly(PR) has a high binding affinity for the tested RNAs, as expected due to its positive charge. Interestingly, this binding shows some sequence specificity, as a higher affinity was observed with the GAAGA motif that was most enriched in the iiCLIP experiment.

**Figure 3.**
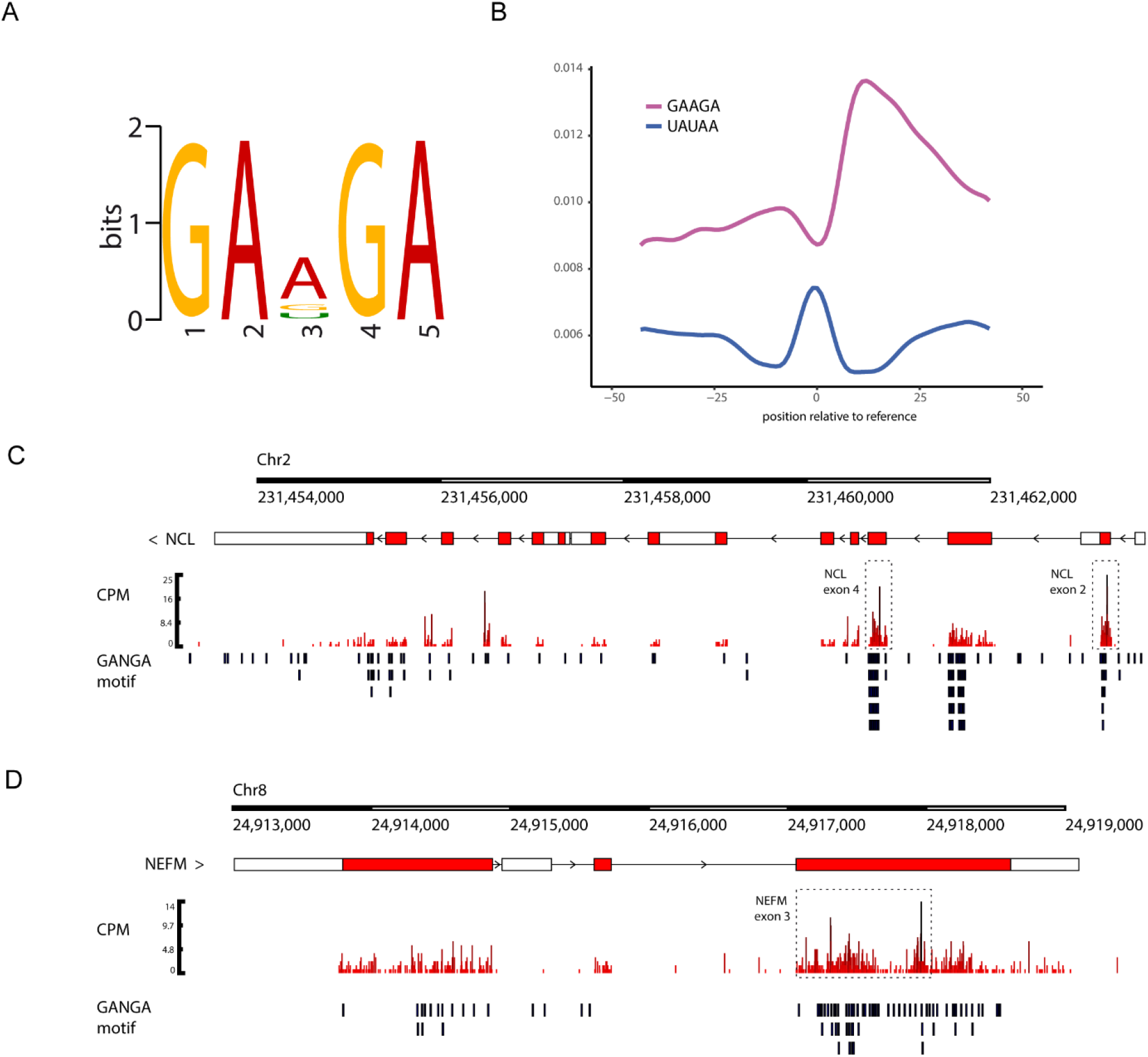
Motif enrichment analysis of poly(PR)-RNA binding. (A) Motif enrichment analysis of the PR100-binding crosslinking sites revealed the most frequent pentamer bound by PR100 was GAAGA (p = 3.1 e^-930^). (B) Analysis of position of the pentamer relative to the crosslinking site. GAAGA is enriched upstream and downstream of the crosslinking site. The UAUAA pentamer has a lower frequency both upstream and downstream of the crosslinking site. (C and D) *NCL* and *NEFM*, which are transcripts highly bound by PR100 in the iiCLIP dataset (Table S1), have frequent GANGA (GAAGA, GAGGA, or GACGA) motifs in proximity to PR100 binding sites. The lower part of each panel indicates the position of these motifs relative to the crosslinking sites within the gene. Gene tracks were normalised by counts per million (CPM).

**Table 1.**
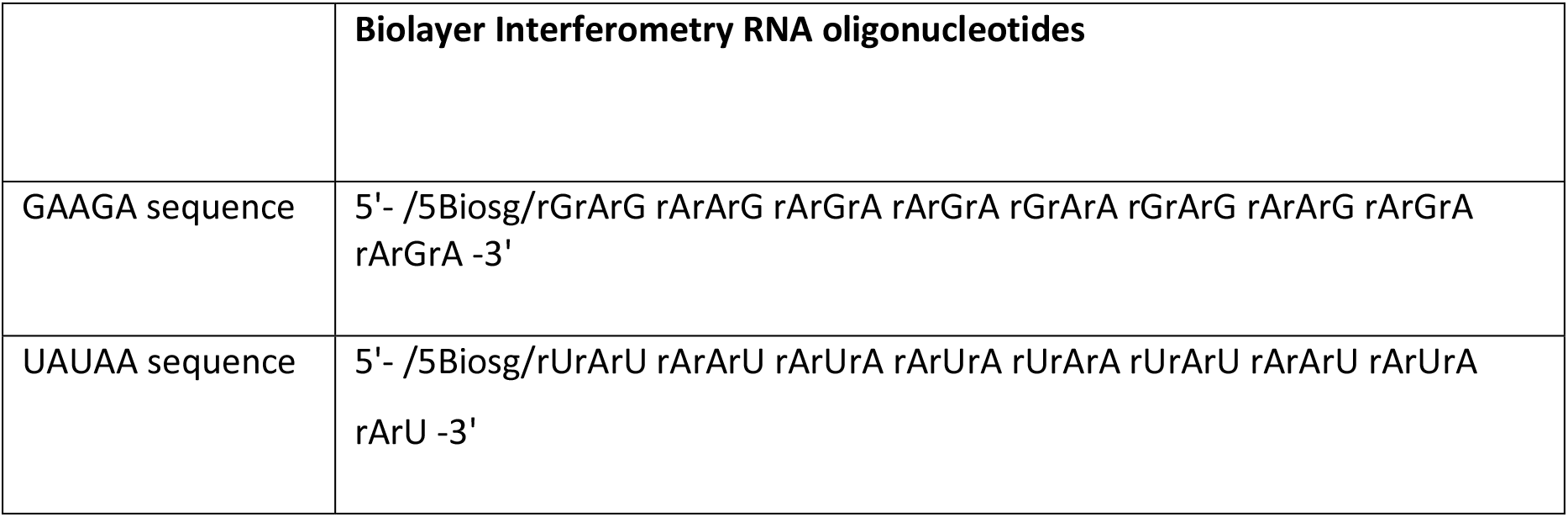
Oligonucleotide sequences.

|  | **Biolayer Interferometry RNA oligonucleotides** |
| --- | --- |
| GAAGA sequence | 5'- /5Biosg/rGrArG rArArG rArGrA rArGrA rGrArA rGrArG rArArG rArGrA rArGrA -3' |
| UAUAA sequence | 5'- /5Biosg/rUrArU rArArU rArUrA rArUrA rUrArA rUrArU rArArU rArUrA rArU -3' |

### PolyGAAGA RNA enhances poly(PR) LLPS

Due to the high affinity of the polyGAAGA RNA sequence to poly(PR), and its enrichment in poly(PR)-binding sites *in vivo*, we investigated whether the polyGAAGA RNA could influence poly(PR) phase separation. Poly(PR) undergoes LLPS in the presence of polyanions, such as RNA (Boeynaems et al., 2017, Boeynaems et al., 2019, Hutten et al., 2020). Thus, we examined whether polyGAAGA RNA had an effect on poly(PR) LLPS. In the absence of RNA, poly(PR) does not undergo LLPS (Fig 5A; top left). By contrast, the addition of bulk HeLa RNA induced the formation of sparse, opaque poly(PR) condensates, which could have irregular morphology (Fig 5A; top right). Remarkably, the presence of equimolar polyGAAGA significantly increased poly(PR) LLPS, indicated by a marked increase in turbidity, and yielded numerous, small round, translucent condensates (Fig 5A; bottom left, and 5B). By contrast, the polyUAUAA RNA induced fewer and larger poly(PR) condensates, which were morphologically distinct from those formed in the presence of polyGAAGA RNA or bulk HeLa RNA (Fig 5A; bottom right). However, polyUAUAA RNA induced less LLPS than polyGAAGA RNA (Fig 5B). These findings suggest that higher affinity RNA, like polyGAAGA, displays enhanced ability to induce poly(PR) condensation.

**Figure 4.**
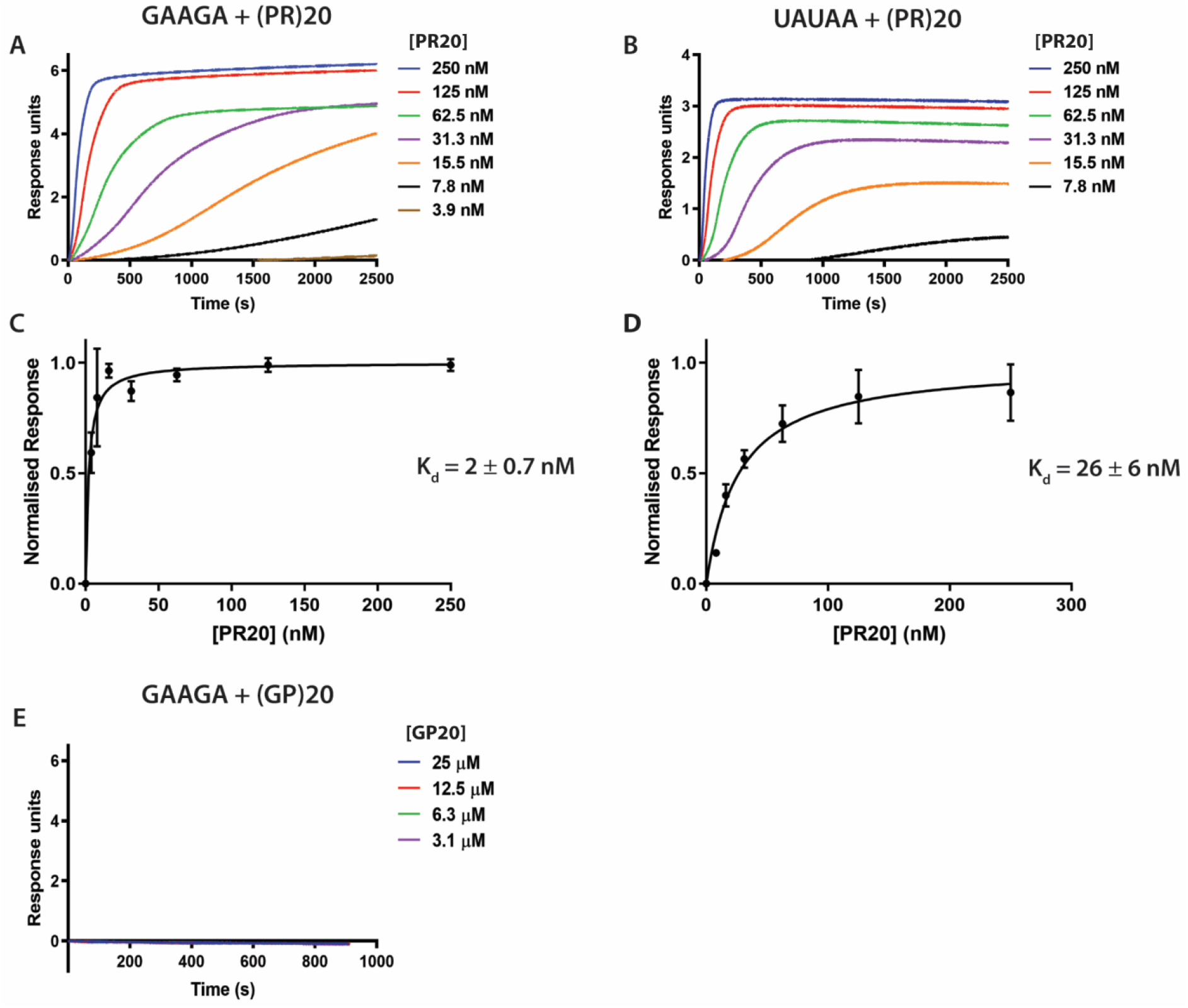
Poly(PR) directly binds to RNA with nanomolar affinity. Biolayer interferometry experiments measuring binding of RNA to PR20. (A-B) The association phases of individual representative experiments. Only PR20 concentrations with responses are shown on the graph. As dissociation was extremely slow (several hours), it was only partially recorded and it is not shown. (C-D) Plots of normalised instrument response versus PR20 concentration. Data represents the average and SD of three independent replicate experiments. (E) Biolayer interferometry experiments measuring binding of polyGAAGA to GP20. 100-fold higher concentrations for GP20 (3.1-25 μM) were used compared to PR20 (1.9-250 nM) to confirm there was no interaction between GP20 and polyGAAGA RNA.

**Figure 5.**
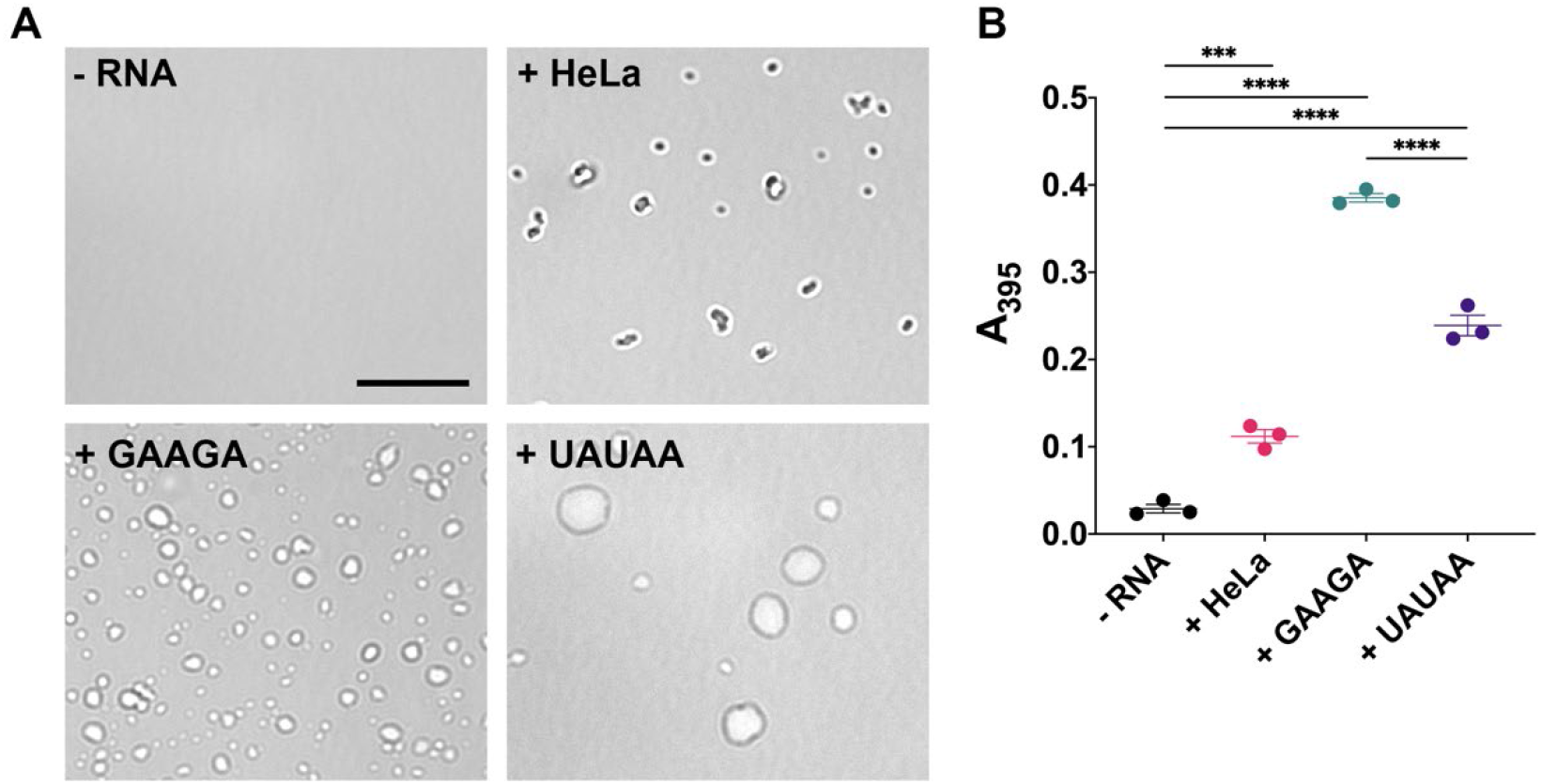
PolyGAAGA RNA enhances poly(PR) LLPS. (A) Condensation of 20 μM PR20-FLAG alone (-RNA), or induced by 20 ng/μL total HeLa cell RNA or equimolar concentrations of either polyGAAGA RNA or polyUAUAA RNA. Representative images from three independent experiments were taken using brightfield microscopy. Black bar represents 10 μm. (B) Turbidity (absorbance at 395nm) measurements of 20 μM PR20-FLAG with or without RNA. Values represent the mean of three independent experiments ± SEM. One-way ANOVA (***p = 0.0003; ****p < 0.0001).

## Discussion

In this study we have investigated whether poly(PR) produced in C9FTD/ALS may directly bind to RNA in human cells. Indeed, we demonstrated direct and specific interactions between the arginine-rich DPR poly(PR) and GAAGA-containing RNAs using a genome-wide approach. Arginine-rich DPRs have been shown to exert deleterious effects on several cellular functions, which include nucleocytoplasmic transport (Zhang et al., 2015, Jovicic et al., 2015, Freibaum et al., 2015, Boeynaems et al., 2016), phase transition of cellular organelles (Lee et al., 2016, Lin et al., 2016, Boeynaems et al., 2017), proteostasis (Kramer et al., 2018) and RNA dysregulation, which has previously been described in C9FTD/ALS models and patient cells and tissue (Yin et al., 2017, Kwon et al., 2014, Kanekura et al., 2016). Some of these effects are likely caused by the interactions of arginine-rich DPRs with other proteins, and our study suggests that their RNA interactions might also contribute to these effects.

Using *in vitro* studies we confirmed that poly(PR) has very strong RNA affinities in the nanomolar range, with a stronger apparent affinity to the polyGAAGA as compared to polyUAUAA RNAs.

Further to the discovery that poly(U) RNA promotes the phase separation of poly(PR) (Boeynaems et al., 2017), it has been shown that in a test tube poly-rA, poly-rU and poly-rC RNA homopolymers can promote phase separation of poly(PR), but poly-rG does so to a lesser extent (Boeynaems et al., 2019). In homopolymeric form, the affinity of interaction between poly(PR) and poly-rA is the strongest, while poly(PR) has almost identical affinity to poly-rU and poly-rC and lowest affinity for poly-rG. This finding has been explained by the ability of poly-rG to form G-quadruplex structures, as opposed to other homopolymeric RNAs, which are unstructured. It has been hypothesised that base stacking interactions associated with G-quadruplex formation could compete with the poly(PR) interaction. Intriguingly, in comparison to these previous findings, the pentameric sequence we found to be most enriched binding to poly(PR) genome-wide *in vivo* has a higher G-content (but with insufficient guanines to form G-quadruplexes) than the least frequently bound pentamer, suggesting that RNA sequences containing guanine can have a high affinity for poly(PR) in the cellular context. Furthermore, it was demonstrated that mixing homopolymeric RNA molecules which can make complementary base pairs, can change the interactions between RNA and poly(PR), possibly due to competition between RNA base pairing interactions and RNA-peptide interactions (Boeynaems et al., 2019). In the RNA sequences we investigated, base pairing is possible in the polyUAUAA but not in polyGAAGA sequence, which may influence the affinity of interaction between poly(PR) and these RNA molecules. It is likely that secondary structures of the RNA influence the interactions determined both *in vivo* and *in vitro*, and the effect of these should be elucidated in further work. Of importance, adding total HEK cell RNA dose dependently ameliorates a nuclear import phenotype induced by adding poly(GR) and poly(PR) to cells (Hayes et al., 2020), suggesting RNA may reduce these phenotypes through high affinity interactions with these DPRs. Intriguingly, we found that RNA sequences that tightly bind to poly(PR) with high affinity have an increased ability to promote poly(PR) condensate formation. It would be of interest in future studies to determine whether these RNA-induced condensates are less toxic to cells. One appealing possibility would be to utilize PR-specific RNA sequences, like polyGAAGA, as “baits” to safeguard the cell by sequestering poly(PR) from deleterious interactions. In fact, TNPO1, a nuclear-import receptor, has been shown to play such a protective role against DPRs when overexpressed (Hutten et al., 2020).

While it was our intention to provide a genome-wide dataset rather than to focus on specific transcripts bound by poly(PR), we report that several interesting RNAs are bound. These include the previously identified paraspeckle long non-coding RNA *NEAT1*, for which polyPR binding was shown to lead to *NEAT1* upregulation (Suzuki et al., 2019) and *NCL*, which encodes the nucleolin protein, which is known to have a more dispersed nuclear localisation in *C9orf72* human tissue and disease models (Haeusler et al., 2014). It is increasingly recognised that RNA dysregulation plays a major role in ALS/FTD and many genetic causes of ALS/FTD are in RNA binding proteins (Nussbacher et al., 2019). Our dataset of poly(PR)-RNA binding can now be used for hypothesis-driven investigation of poly(PR) effects on RNAs. Understanding the biology of these interactions may help to further elucidate the underlying aetiology of neurodegeneration in C9FTD/ALS.

## Materials and Methods

### Cell lines

HEK293Ts were cultured in Dulbecco’s modified Eagle medium supplemented with 10% fetal bovine serum, grown at 37°C with 5% CO2 injection, and routinely passaged.

### Transient transfections and iiCLIP protocol

PR100 and GA100 (Mizielinska et al., 2014) were cloned into pcDNA5 Flp-In Expression vectors with a 3’ triple FLAG tag, generating PR100-3xFLAG (PR100-FLAG) and GA100-3xFLAG (GA100-FLAG) pcDNA5 plasmids, with the 3xFLAG only vector (FLAG) also used as a control. For iiCLIP experiments, HEK293Ts were grown at ≈80% confluency in 10 cm plates and transiently transfected with PR100-FLAG, GA100-FLAG or FLAG pcDNA5 plasmids using Lipofectamine 2000. Expression of the constructs was induced by supplementing the media with 150 ng ml^-1^ of doxycycline for 24 hours.

The iiCLIP protocol was performed as previously described (Lee et al., 2021). Transiently transfected cells induced for 24 hours were irradiated with UV once with 160 mJ/cm2 using Stratalinker 1800 at 254 nm. Protein-RNA complexes were ligated to a preadenylated, infrared dye labeled adaptor and purified. RNA was isolated using proteinase K digestion and reverse-transcribed into cDNA. cDNA was subsequently purified and circularised.

### iiCLIP analysis

Multiplexed cDNA libraries were sequenced using Illumina HiSeq, generating 100-nt single-end reads. Sequenced reads were processed by the iMaps software package (http://icount.biolab.si/), and de-multiplexed into individual libraries based on their experimental barcodes. Unique molecular identifier nucleotides were used to distinguish and collapse PCR duplicates. The barcode sequences and adaptors were removed from the 5’ and 3’ ends. Trimmed sequences were mapped to the human genome (build GRCh38, Gencode version 27) with STAR aligner allowing two mismatches (Langmead and Salzberg, 2012). Uniquely mapping reads were kept, and the preceding aligned nucleotide was assigned as the DPR crosslinked site. Significant crosslink sites were determined by the iCount False Discovery Rate (<0.05) algorithm by weighing the enrichment of crosslinks versus shuffled random positions (https://github.com/tomazc/iCount). For subsequent analysis, we set a threshold of at least 300,000 unique crosslink events for each PR100-crosslinked sample, and n=4 samples met this threshold. For three of these samples, protein-RNA complexes had been purified using SDS-PAGE and transferred onto nitrocellulose membranes, as previously described (Lee et al., 2021). For one of these samples, protein-RNA complexes had been purified using immunoprecipitation with beads. We analysed the number of crosslinking events for these PR100-crosslinked samples and their corresponding controls: PR100-non-crosslinked, GA100-crosslinked, GA100-non-crosslinked and FLAG-crosslinked. Crosslinking events were normalised by the total number of crosslinks in the sample per million (Counts per million - CPM). For gene level analysis, genes were identified that contain at least 200 crosslink events from PR100-FLAG iiCLIP, with <10% binding in the control conditions of FLAG crosslinked samples and PR100-FLAG non-crosslinked samples (Table S1). These genes were used for ontology enrichment analysis performed with R package ClusterProfiler, comparing against all other genes, using an FDR correction and adjusted p value cut off of <0.01 (Yu et al., 2012). Enriched pentamers were calculated with Dreme v.5.4.1 (Bailey, 2011) using the 5 nt upstream and 30 nt downstream to the significant crosslink sites, compared to similar sequences collected from random positions of the same genes that did not overlap with any significant crosslink sites.

### Biolayer Interferometry Measurements

Biolayer Interferometry experiments were performed on a ForteBio Octet Red 96 instrument (Sartorius). Biotinylated RNAs were synthesised by Integrated DNA Technologies. PR20 and GP20 peptides were synthesised by Cambridge Research Biochemicals. Biotinylated RNA and poly-DPR peptides were dissolved in 0.22 μm filtered Tris-EDTA (10 mM Tris-HCl, 1 mM disodium EDTA, pH 8.0) buffer solution with 150 mM NaCl, 0.1 mg/mL bovine serum albumin (BSA) and 0.01% Tween-20 to reduce non-specific interactions. The assays were carried out at 25°C in a 96-well plate and a sample volume of 200 μL. Streptavidin-coated biosensors were pre-equilibrated, loaded with biotinylated RNAs, and exposed to protein concentrations ranging from 1.9 – 250 nM for PR20 and 3.1-25 μM for GP20. Equilibrium dissociation constants (K_d_) for the RNA–protein interactions were determined by plotting the instrument response at equilibrium as a function of protein concentration and fitting the data assuming a 1:1 interaction, using non-linear least squares regression using an in-house software. Oligonucleotide sequences are provided in Table 1. Biological triplicates were performed using freshly prepared RNA and protein solutions in independent experiments.

### *In vitro* poly(PR) condensation assay

Poly(PR) 20-mer dipeptide repeat protein with a C-terminal FLAG tag was purchased from CSBio and verified by mass spectrometry. Sequence of poly(PR): PRPRPRPRPRPRPRPRPRPRPRPRPRPRPRPRPRPRPRPRGSFEGDYKDDDDK. Lyophilized powder was reconstituted in 1X PBS and snap frozen in single use 200 μM aliquots and stored at −80°C. PolyGAAGA and polyUAUAA RNA sequences were ordered from IDT (sequences provided in Table 1). Lyophilized powder was reconstituted in RNase free water, to a final stock concentration of 100 μM. Aliquots were snap frozen and stored at −20°C. For RNA-induced condensate formation, poly(PR) and all RNAs were first thawed on ice. Poly(PR) was diluted to a final concentration of 20 μM in 20 mM HEPES-NaOH (pH 7.4), 150 mM NaCl and 1mM DTT. HeLa RNA at a final concentration of 20 ng/μL or equimolar amounts (20 μM) of either polyGAAGA or polyUAUAA RNA were added to poly(PR) and incubated at room temperature for 30 minutes. Phase-separated condensates were then imaged by Brightfield microscopy (EVOS M5000). For turbidity measurements, absorbance values were read at an absorbance of 395 nm using TECAN (Safire^2^). Statistical analyses were performed using GraphPad Prism 8. Details are given in the figure legend.

---

**Figure S1.**
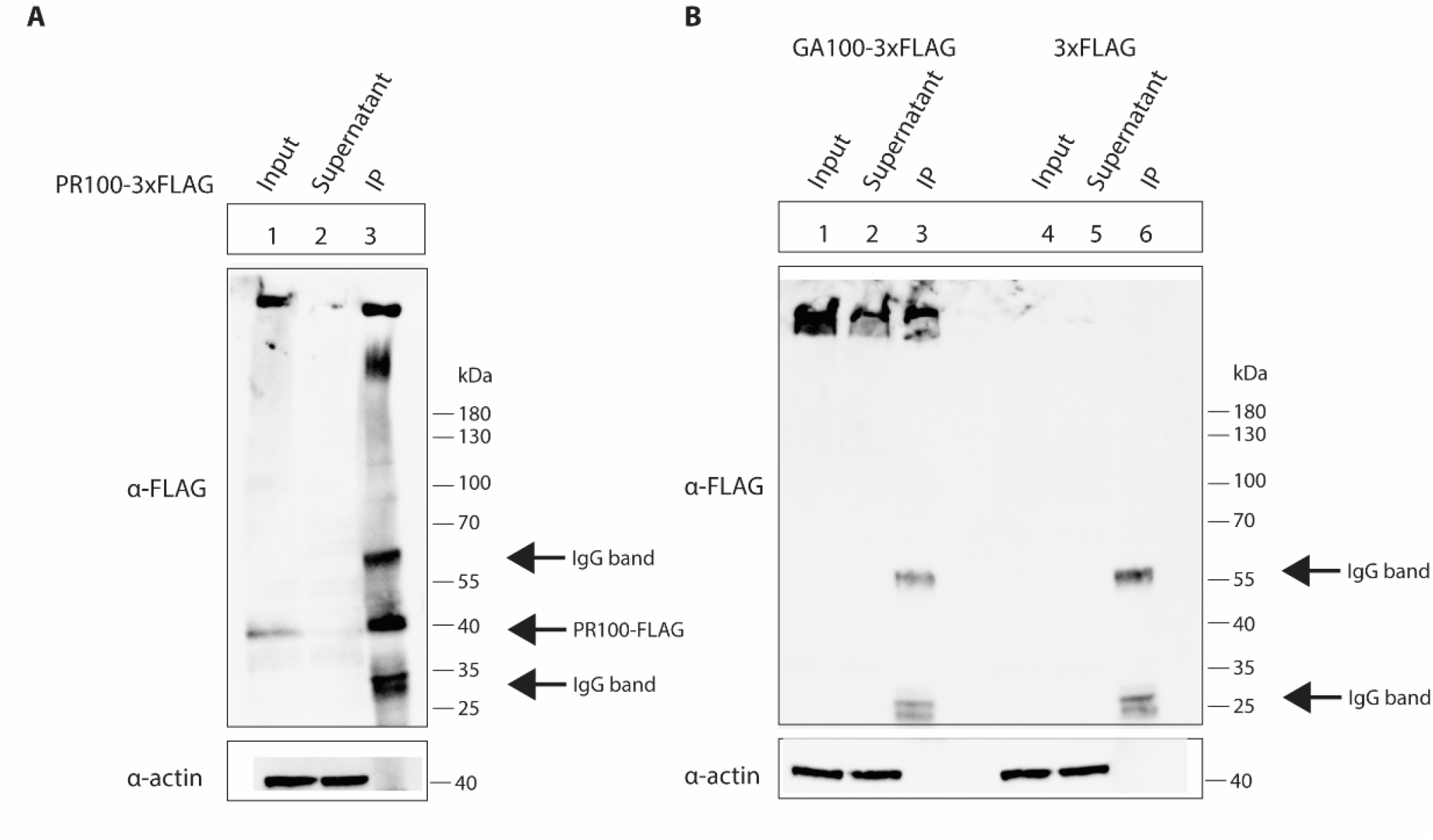
Immunoprecipitation of FLAG-tagged constructs. (A-B) HEK293Ts were induced to express PR100-FLAG, GA100-FLAG or FLAG for 24 hours, then UV crosslinked. Protein-RNA complexes were immunoprecipitated using the FLAG tag. PR100-FLAG, GA100-FLAG and actin are detected in the cell lysates, PR100-FLAG and GA100-FLAG are detected in the immunoprecipitates, whilst actin is not detected.

**Figure S2.**
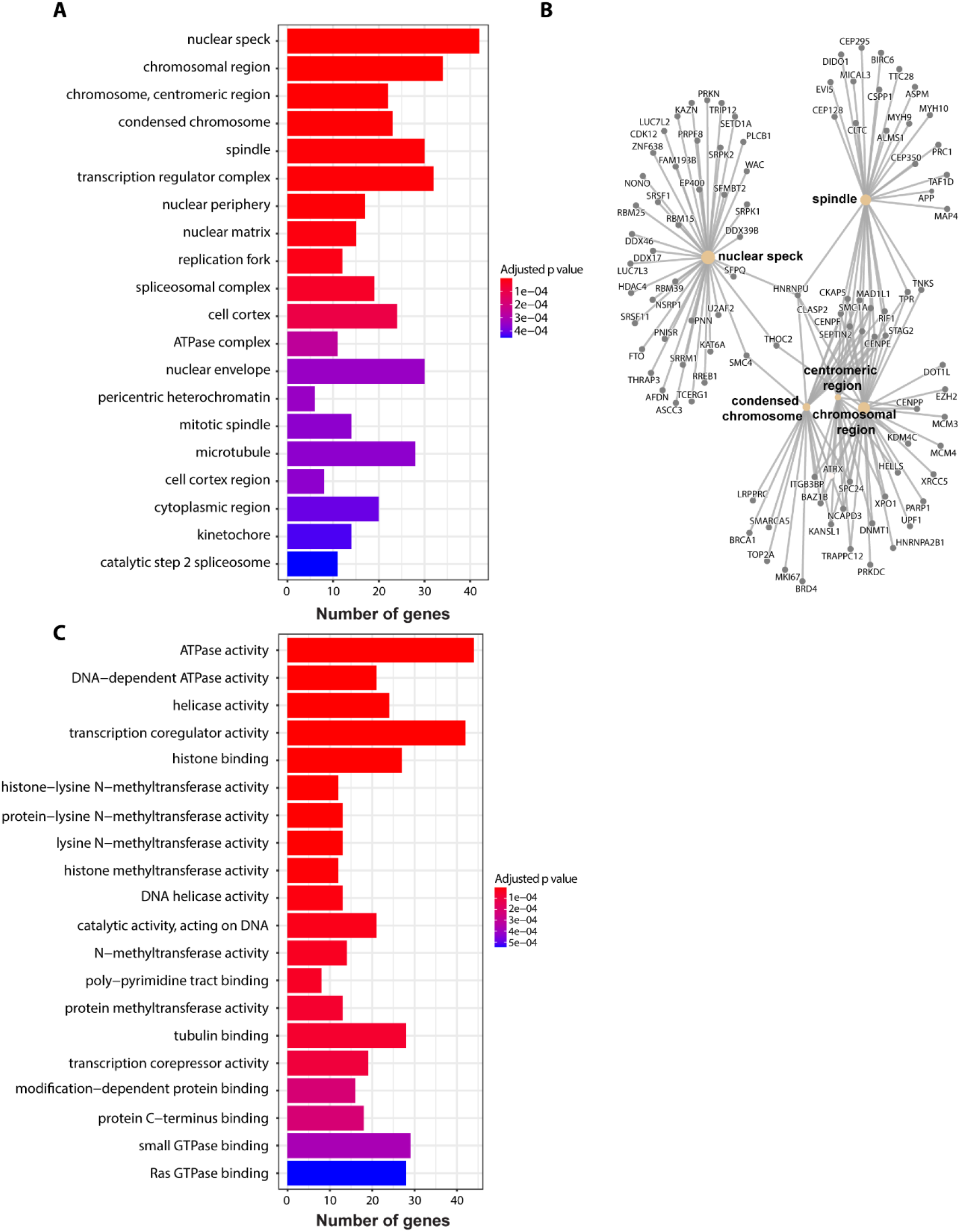
Gene Enrichment Analysis of PR100 crosslinked RNAs. Gene Ontology gene set enrichment analysis of RNAs bound in PR100 samples are represented by their Cellular Component (A) and their Molecular function (C). The number of genes from the PR100-FLAG crosslinking dataset in each Gene Ontology category are shown and colour coded by adjusted p value. (B) The genes of RNAs bound in PR100 samples from the top three significant categories within Cellular Component (nuclear speckles, chromosome region and centromeric chromosome regions) are represented in a gene-concept network. The size of the circle for each Cellular Component is proportional to the number of genes identified within that category.

